# Gasdermin D mediates a fast transient release of ATP after NLRP3 inflammasome activation before ninjurin 1–induced lytic cell death

**DOI:** 10.1101/2024.12.10.627750

**Authors:** Julieta Schachter, Adriana Guijarro, Diego Angosto-Bazarra, Miriam Pinilla, Laura Hurtado-Navarro, Etienne Meunier, Ana B. Pérez-Oliva, Pablo J Schwarzbaum, Pablo Pelegrin

## Abstract

Pyroptosis is a lytic cell death triggered by the cleavage of gasdermin (GSDM) proteins and subsequent pore formation by the N-terminal domain oligomerization in the plasma membrane. GSDMD is cleaved by caspase-1/-4/-5/-11 upon inflammasome activation and mediates IL-1β and IL-18 release. GSDMD pores favors ninjurin-1 (NINJ1) induced plasma membrane rupture and cell death. Here, we demonstrate that GSDMD mediates early ATP release upon NLRP3 inflammasome activation independently of NINJ1, occurring before IL-1β release and cell death and constituting an early danger signal. The release of ATP is a transient signal terminated before the cells continue to permeabilize and die. The different N-terminal of GSDMA to E are also able to release ATP and induce monocyte migration towards pyroptotic cells. This study reveals ATP release as an early, and transient danger signal depending on GSDMD plasma membrane permeabilization, independently of the late stages of lytic cell death.

## INTRODUCTION

In the extracellular space, adenosine triphosphate (ATP) functions as a signaling molecule, engaging purinergic receptors and initiating different signaling cascades involved in various physiological processes, with important roles in neurotransmission, cell death, vasodilatation or cell differentiation^1^. In particular, extracellular ATP (eATP) is widely recognized as a danger signal and plays a crucial role in modulating several steps of the immune response. Extensive research has demonstrated the impact of eATP and/or some of its metabolic byproducts on inflammation, chemotaxis, antigen presentation, modulation of T cell function, and regulation of macrophage activity^2,3,4^. Several stimuli such as hypoxia, shear stress, swelling or the presence of pathogens can trigger ATP release through channels, pores and exocytosis^5,6,7^. Once in the extracellular milieu, ATP is hydrolyzed by ecto-nucleotidases, a family of membrane-bound enzymes with catalytic sites facing the extracellular space^8^. Ecto-nucleotidase activity not only terminates the action of eATP on specific purinergic receptors, but also generates different ATP byproducts in the extracellular space capable of signaling to different purinergic receptors^9^.

Inflammation can be initiated by engaging the purinergic receptor P2X subunit 7 (P2X7), which responds to mM concentrations of eATP. P2X7 receptor mediates the efflux of intracellular K^+^ and the subsequent activation of the nucleotide-binding domain and leucine-rich repeat-containing receptor with a pyrin domain 3 (NLRP3) inflammasome^10,11^. In turn, NLRP3 inflammasome leads to caspase-1 activation and the cleavage of different cellular proteins, including pro-interleukin (IL)-1β and pro-IL-18, as well as gasdermin D (GSDMD)^12^. The resulting GSDMD N-terminal fragment is separated from its C-terminal repressor domain and oligomerizes in the plasma membrane, forming pores that allow the direct release of IL-1β and IL-18^13,14^. GSDMD-pores in the plasma membrane favor the oligomerization of the protein ninjurin 1 (NINJ1) that induces large ruptures in the membrane, leading to the lytic cell death termed pyroptosis^15,16,17^. Pyroptosis is a highly inflammatory form of cell death characterized by cell swelling, membrane rupture, and the release of intracellular content, including different intracellular proteins, cytokines, inflammasome oligomers and mitochondrial DNA^18,19,20^.

While the effects of eATP on NLRP3 inflammasome activation are well documented, the inflammatory signals promoting ATP release are less known. In this study we found that, following NLRP3 inflammasome activation, ATP is released from macrophages as an early and transient danger signal, causing a non-linear pattern of eATP accumulation dynamically controlled by ecto-nucleotidase activity. We demonstrate that ATP release is a regulated process that requires GSDMD membrane permeabilization and precedes IL-1β release and cell death. The N-terminal fragments of GSDMA, B, C, D and E were all also able to induce ATP release, with the accumulated eATP inducing monocyte migration. Furthermore, the transient increase in eATP induced by NLRP3 activation disappears before the pyroptotic lytic cell death process begins, showing that the late phases of NINJ1–mediated pyroptotic lytic cell death occurs without ATP release.

## RESULTS

### NLRP3 inflammasome activation induces ATP release by GSDMD

Activation of the NLRP3 inflammasome by nigericin in LPS-primed bone marrow derived macrophages (BMDMs) triggered a dose-dependent ATP release, causing a time-dependent increase of eATP concentration, followed by rapid degradation (Fig. 1A). A similar eATP kinetic profile was observed when the same treatment was applied to macrophages from P2X7 receptor deficient mice (Fig. 1B, F), thus implying that this receptor is not required for the response. ATP release was inhibited by the specific NLRP3 inhibitor MCC950 (Fig. 1A, B), and was not observed in resting unprimed macrophages (Fig. 1C), demonstrating the dependency of NLRP3.

**Figure 1.**
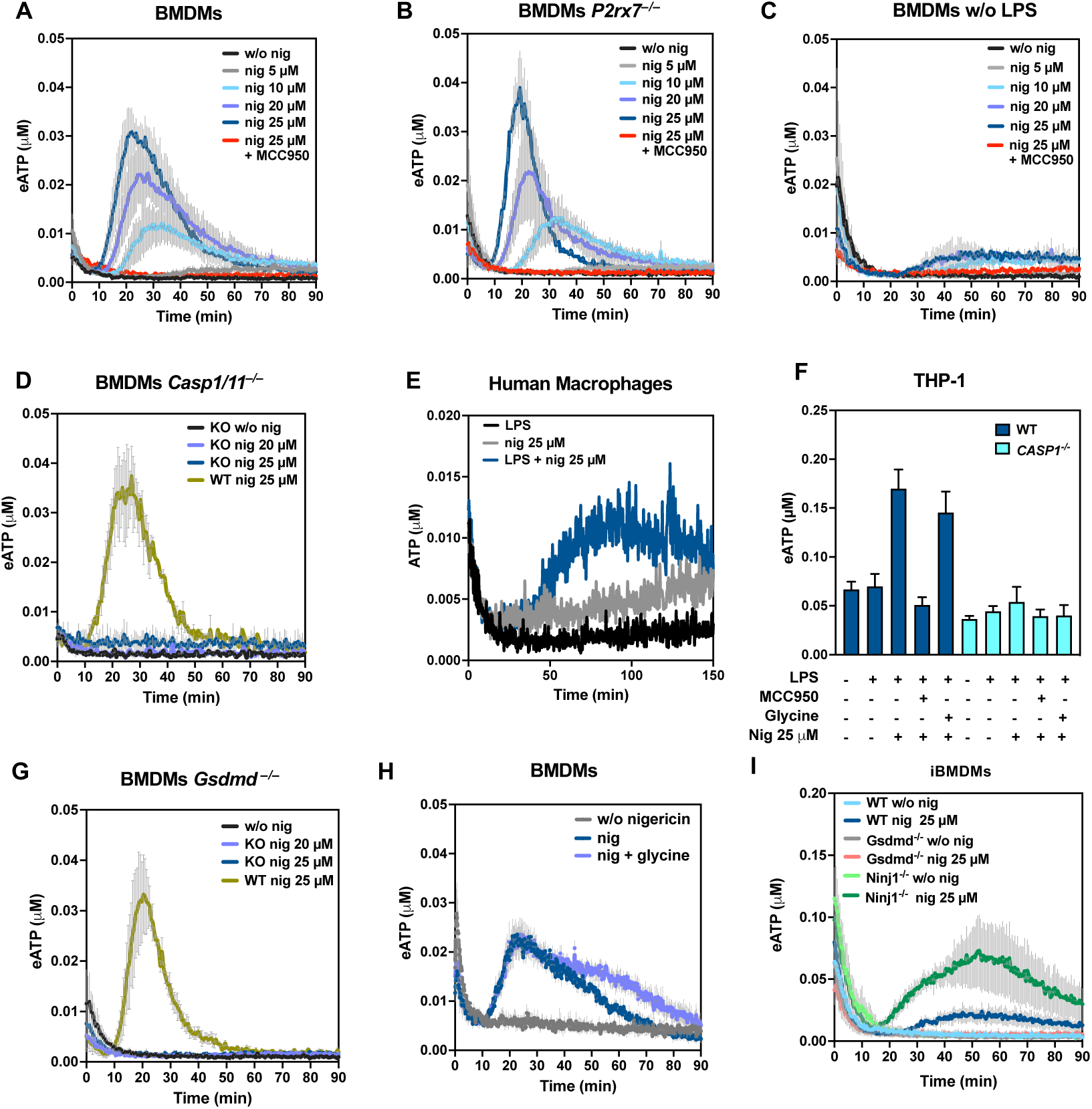
ATP release after NLRP3 activation is GSDMD–dependent. **A-E**. Kinetics of extracellular ATP (eATP) after canonical NLRP3 inflammasome activation in LPS–primed mouse bone marrow derived macrophages (BMDMs) (A, B, D) or unprimed BMDMs (w/o LPS) (C), obtained from wild type (A, C), *P2rx7*^−/−^ (B) or *Casp1/11*^−/−^ (D) mice. The different curves show eATP concentration of macrophages treated with saline vehicle solution (black lines, w/o nig), nigericin (nig) 5 µM (grey lines), 10 µM (light blue lines), 20 µM (blue lines), 25 µM (dark blue lines) or nigericin 25 µM with MCC950 10 µM (red lines). To compare results using BMDMs from wild type (WT) mice *vs* the knock-out (KO) genotype, the green lines in D represent eATP kinetics of BMDMs wild type treated with nigericin 25 µM. Data are expressed in eATP concentration (µM) as the mean ± SEM of independent experiments (N) performed in duplicate (n), with N = 3, n = 2 for cells from wild type mice, and N = 2, n = 2 for cells from *P2rx7*^−/−^ and *Casp1/11*^−/−^ mice. **E.** Kinetics of eATP from LPS–primed human monocyte–derived macrophages treated with saline solution (black line), nigericin 25 µM (blue line), or in unprimed macrophages treated with nigericin 25 µM (grey line). Representative experiment of three experiments performed in duplicate are shown. **F.** THP-1 WT cells (dark blue bars) and THP-1 *CASP1*^−/−^ (light blue bars) cells were primed or not with LPS and treated or not with nigericin 25 µM, MCC950 10 µM or glycine 5 mM, as indicated in the bar graph. Data are expressed in eATP concentration (µM) at 120 minutes as the mean ± SEM of N = 3, n = 2. **G.** Kinetics of eATP of BMDMs from wild type (green line) or *Gsdmd*^−/−^ (black, light blue and blue lines) mice treated as in A. Data are expressed in eATP concentration (µM) as the mean ± SEM of N = 3, n = 2. **H.** LPS–primed BMDMs were treated with saline vehicle solution (grey line), nigericin 25 µM (blue line) or nigericin 25 µM with glycine 5 mM (light blue line). Data are expressed in eATP concentration (µM) as the mean ± SEM of N = 3, n = 2. **I.** LPS–primed immortalized BMDMs WT, *Gsdmd*^−/−^ or *Ninj1*^−/−^ treated with saline solution (light blue for WT, grey for *Gsdmd*^−/−^ and light green for *Ninj1*^−/−^) or nigericin 25 µM (dark blue for WT, red for *Gsdmd*^−/−^ and dark green for *Ninj1*^−/−^). Data are expressed in eATP concentration (µM) as the mean ± SEM of N = 3, n =2. See also Figures S1, S2 and S3.

Since NLRP3 activation in macrophages leads to caspase-1 dependent cell death, we next analyzed ATP release in *Casp1/11^−/^*^−^ BMDMs. We found that ATP release was absent when *Casp1/11^−/^*^−^ BMDMs were primed with LPS and treated with nigericin (Fig. 1D).

We also tested the capacity for ATP release from human macrophages derived from blood monocytes (Fig. 1E) as well as from THP-1 and *CASP1^−/^*^−^ THP-1 cells (Fig. 1F) after NLRP3 activation with LPS and nigericin, in the presence or absence of MCC950. We confirmed that LPS–primed human macrophages were able to release ATP after nigericin treatment and this release was dependent on NLRP3 and caspase-1.

BMDMs from mice deficient in GSDMD failed to accumulate eATP after NLRP3 inflammasome activation (Fig. 1G), indicating that GSDMD induced plasma membrane permeabilization is required to release ATP. Since lytic cell death during pyroptosis downstream of GSDMD-plasma membrane permeabilization is driven by NINJ1, we investigated whether the ATP release associated with GSDMD activity occurs through NINJ1–dependent plasma membrane rupture. Accordingly, we measured the eATP kinetics in nigericin-treated, LPS-primed BMDMs in the absence or presence of glycine, which inhibits NINJ1 membrane clustering and avoids plasma membrane rupture^13,21,22^. We found that glycine did not affect ATP release (Fig. 1F, H) induced by nigericin, suggesting that NINJ1 is not involved in the ATP release during pyroptosis. As a control, we found that glycine blocked LDH release, an estimator of membrane damage, but not IL-1β release after NLRP3 activation (Fig. S1A). We confirmed that NINJ1 is not involved in ATP release after NLRP3 activation by using immortalized BMDMs *Ninj1*^−/−^. These cells, treated with LPS and nigericin, exhibited ATP release, while, as expected, immortalized BMDMs *Gsdmd*^−/−^ did not (Fig. 1I). Accordingly, immortalized BMDMs *Ninj1*^−/−^, as well as *Gsdmd*^−/−^, released very low amounts of LDH compared with wild type macrophages in response to the same treatment (Fig. S1B–D).

During treatments that induced ATP release, an early phase of eATP decrease was observed, though it was not included in the analysis. This decrease, likely due to mechanical stimulation, was independent of NLRP3 activation and pyroptosis, as it was unaffected by the NLRP3 inhibitor MCC950 and was not observed in *Gsdmd*^−/−^ macrophages (Fig. 1A-C, G-I).

Although pannexin-1 and connexin-43 are reported to mediate ATP release in macrophages^23^, carbenoxolone (CBX) and gadolinium (Gd^3+^), two generic inhibitors of these and other ATP conduits, did not affect eATP kinetics (Fig. S2A,B). This is in line with the idea that during NLRP3 induced pyroptosis, ATP might be transported to the extracellular milieu through GSDMD pores. Overall, these results show that ATP release in macrophages is triggered soon after the activation of the NLRP3 inflammasome, following a process requiring caspase-1 activity and the presence of GSDMD. This process is not linked to the late phases of pyroptosis mediated by NINJ1 plasma membrane lysis.

### Extracellular ATP is hydrolyzed on the surface of macrophages during early pyroptosis

In the extracellular milieu, eATP can be hydrolyzed to extracellular ADP (eADP) and inorganic phosphate by ecto-nucleotidases. To evaluate the ecto-ATPase activity present on the surface of LPS-primed macrophages after treatment with nigericin, we added increased concentrations of exogenous ATP to the assay medium, with these concentrations falling within the same range as the endogenous eATP concentrations shown in Fig. 1.

ATP addition led to an immediate increase in eATP to a maximum, which allowed us to determine the subsequent eATP decrease kinetics (Fig. S3A). The initial lineal decay in eATP was used to estimate ecto-ATPase activity at each ATP concentration (Fig. S3B). The ecto-ATPase activity of the macrophages followed a linear function with ATP concentration, with the slope of the curve (K_ATP_) amounting to 3.2 μM eATP/μM eATP (min.mg of protein) (Fig. S3B). As compared to ecto-ATPase activity of other cell types^24^, BMDMs presented a relatively high eATP hydrolysis rate. Thus, the observed fast eATP degradation during pyroptosis is expected to play a regulatory role, preventing excessive and continuous ATP signaling.

### ATP release dependent on NLRP3 activation precedes lytic cell death

To gain further insights into the relation of ATP release and cell death, we studied the kinetics of YO-PRO-1 (629 Da) uptake in LPS-primed BMDMs treated with nigericin. The average behavior of BMDMs show that YO-PRO-1 uptake displays a sigmoidal internalization kinetics, with an initial lag phase of basal fluorescence, followed by a rapid increase in dye uptake. Such a response was completely absent in *Gsdmd*^−/−^ macrophages or in cells treated with MCC950 (Fig. 2A). An analysis of individual cells shows that, following a lag phase of different magnitudes, the sigmoidal phenomenon was sufficiently steep to be described as a near “all or none” uptake process (Fig. 2B).

**Figure 2.**
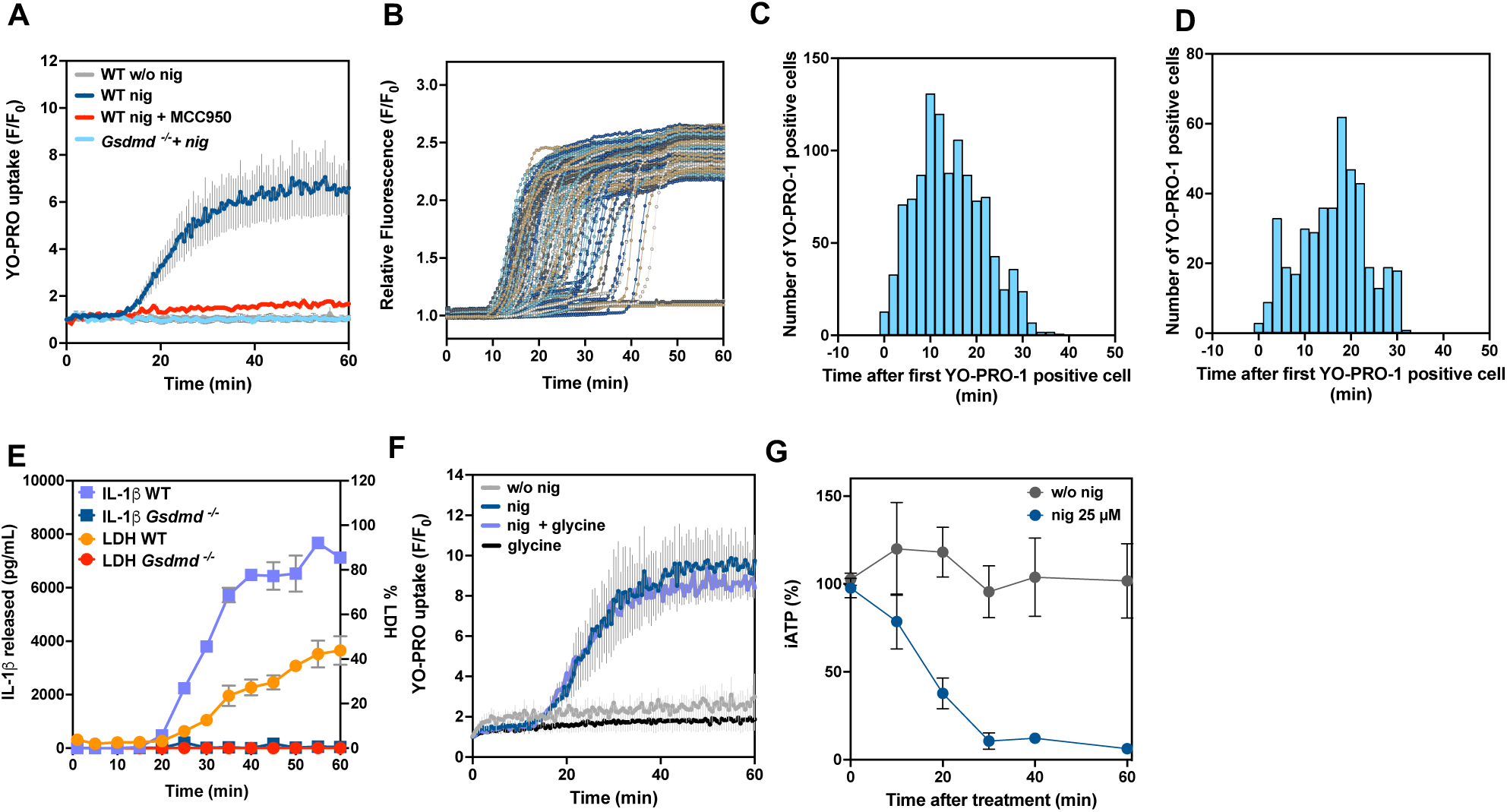
ATP release precedes pyroptotic lytic cell death. **A.** YO-PRO-1 uptake from LPS-primed BDMDs from WT mice treated with nigericin (dark blue line), or nigericin (nig) with 10 µM MCC950 (red line) or from unprimed cells treated with nigericin (grey line) or from LPS-primed BMDMs from *Gsdmd*^−/−^ mice treated with nigericin (light blue line). Data are the mean ± SEM of independent experiments (N) performed in duplicate (n), N = 3, n = 2, and the fluorescence was normalized to the initial florescence in each experiment (F/F_0_). **B.** YO-PRO-1 uptake of individual LPS-primed BMDMs treated with 25 µM nigericin at time = zero (each macrophage is represented as a line). Representative experiment of 5 independent experiments. **C,D**. Histogram showing the number of YO-PRO-1 positive cells obtained from LPS-primed BMDMs from wild type (WT) (C) or *P2rx7*^−/−^ (D) mice at different times of nigericin application. Each experiment was normalized by setting the time of the first YO-PRO positive cell as time 0. Total number of cells = 1098 (WT) and 432 (*P2rx7*^−/−^), obtained from 5 (WT) or 3 (*P2rx7*^−/−^) independent experiments, with approx. 200 cells quantified in each experiment. **E.** Kinetic of IL-1β and LDH release from LPS-primed BDMDs treated with 20 µM nigericin. Light blue and orange lines from WT mice; or red and blue lines from *Gsdmd*^−/−^ mice. Results are the mean ± SEM for *N* = 3, n = 2. **F.** YO-PRO-1 uptake from LPS-primed BDMDs treated with saline solution (grey line), 5 mM glycine (black line), 25 µM nigericin (dark blue line) or 25 µM nigericin and 5 mM glycine (light blue line). Results are the mean ± SEM for *N* = 3, n = 2. **G.** Kinetic of intracellular ATP concentrations after canonical NLRP3 inflammasome activation in LPS-primed BMDMs treated with 25 µM nigericin (blue symbols) or not (grey symbols). Results are expressed as the percentage of total intracellular ATP and are the mean ± SEM for N = 3 independent experiments.

A frequency histogram of YO-PRO-1-positive cells was built, normalizing the initial time point (t = zero) to the moment the first cell in each experiment became YO-PRO positive. This analysis revealed that the majority of the macrophages (80.4 %) exhibited a burst of dye uptake between 0 and 20 min (Fig. 2C), with a peak at 14 minutes. The average time until the first cell became permeabilized was 23.3 ± 0.5 minutes, and after 30 min macrophages that did not capture Yo-Pro-1 remained unpermeabilized through the recording time.

When the same analysis was done in BMDMs from a P2X7 receptor-deficient mice, the histogram shifted to the right, with a peak at 18 minutes and an average time to the first cell’s permeabilization of 31.1 ± 0.5 minutes (Fig. 2D). Moreover, the percentage of permeabilized cells differed significantly, with 85.5 % of YO-PRO-1 positive in WT cells compared to 53.7 % in *P2×7*^-/-^ cells, suggesting a potential function of P2X7 on GSDMD pore formation.

In Fig. 1B we showed that P2X7 receptor is not required for ATP release in response to NLRP3 activation. Therefore, the observed delay in the YO-PRO-1 uptake and the reduced number of positive cells in P2X7-deficient cells are due to the absence of P2X7 receptor. This suggests that the released ATP can stimulate YO-PRO-1 uptake in neighboring cells.

We also compared eATP kinetics with IL-1β release, a phenomenon associated with GSDMD pore formation, and the release of LDH associated to GSDMD–induced NINJ1 mediated cell death. Both, IL-1β and LDH release exhibited non-linear release kinetics upon nigericin stimulation. However, the increase in IL-1β release was significantly faster and greater in relative magnitude compared to LDH (Fig. 2E), suggesting two qualitative different processes.

Glycine did not affect the YO-PRO-1 uptake (Fig. 2F) induced by nigericin, suggesting that it measured GSDMD plasma membrane permeabilization. Therefore, YO-PRO-1 uptake and IL-1β release seem be coincident in time and dependent on GSDMD, LDH release is right-shifted and dependent on NINJ1, and ATP release is the earliest event dependent on the initial GSDMD permeabilization. The fact that YO-PRO-1 uptake appears after ATP release could mean that ATP diffusion thought initial GSDMD pores will be easy or could impair than the influx of YO-PRO-1 or that YO-PRO-1 uptake does not effectively label early or transient GSDMD pores, as transient GSDMD pores are ∼10% the magnitude of a permeabilized cell^25^.

When NINJ1–dependent LDH release initiates, most cells are already GSDMD–permeabilized, and eATP concentration starts to decrease to levels similar to the resting state. This suggests a novel concept, i.e., that the phase of plasma membrane rupture during pyroptosis may not be the primary moment for significant ATP release. In fact, besides the effect of ecto-nucleotidases removing eATP, the intracellular ATP levels also decline over time following NLRP3 activation, dropping to less than 2% after 30 minutes (Fig 2G). This ensures that even when large portions of the plasma membrane are damaged by NINJ1, ATP is no longer being released from cells. Thus, given the observed GSDMD dependence of ATP release without the participation of hemichannels, and the fact that activated GSDMD forms large pores high enough to transport ATP but not LDH, it is tempting to speculate that GSDMD acts as an ATP conduit, allowing ATP release early on, before plasma membrane rupture. By the time LDH is released via NINJ1 oligomerization, intracellular ATP levels have dropped so low that there is insufficient chemical transmembrane gradient to drive further ATP release.

### ATP is released through different gasdermins and ninjurin 1

To corroborate the role of GSDMD pore formation in ATP release, we studied the kinetics of ATP release in a recombinant system of HEK-293T cells expressing either full-length GSDMD or the GSDMD N-terminal domain. Discrete ATP concentrations were measured in the culture medium at different time points post-transfection. Cells transfected with GSDMD N-terminal showed increased eATP levels at all time points, peaking at 6 hours (Fig. 3A).

**Figure 3.**
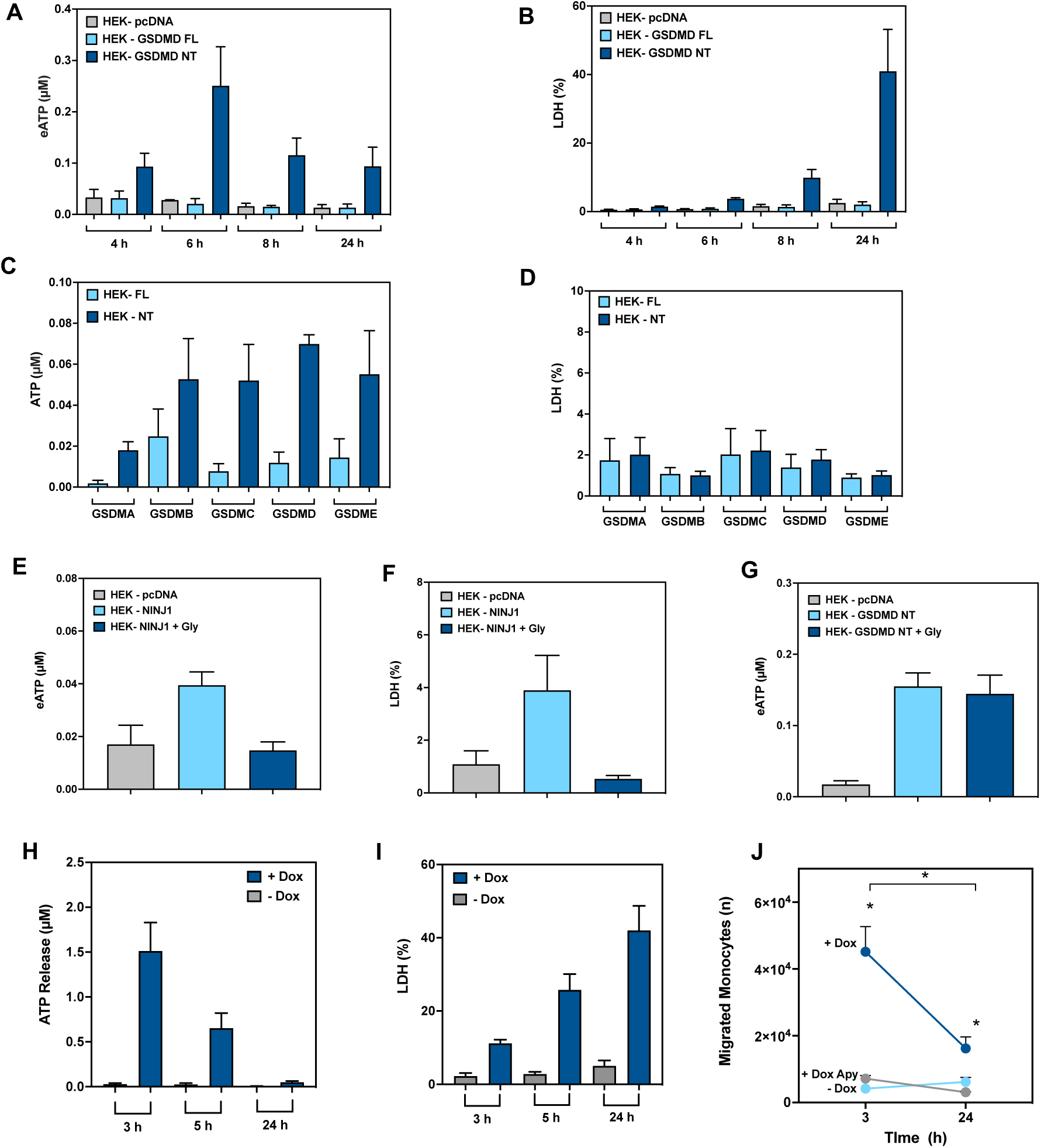
ATP is released from different GSDMs and NINJ1 and promotes cell migration. **A,B**. Extracellular ATP (eATP) (A) and LDH (B) from HEK-293T cells transfected with GSDMD full-length (FL), GSDMD N-terminal (NT) or with pcDNA3.1 at 4, 6, 8 and 24 hours after transfection. Results are the mean ± SEM of independent experiments (N) performed in duplicate (n), N = 4, n = 2. **C,D**. eATP (C) and LDH (D) from HEK-293T cells transfected with GSDMA, GSDMB, GSDMC, GSDMD and GSDME FL and NT at 4 hours after transfection. Results are the mean ± SEM of N = 3, n = 2. **E,F**. eATP (E) and LDH (F) from HEK-293T cells transfected with pcDNA3.1 (control, grey bars), NINJ1 (light blue bars) or NINJ1 with 5 mM glycine (Gly) (dark blue bars) at 6 hours after transfection. Results are the mean ± SEM of N = 4, n = 2. **G**. eATP from HEK-293T cells transfected with pcDNA3.1 (control, grey bars), GSDMD NT (light blue bars) or GSDMD NT with 5 mM glycine (dark blue bars) at 8 hours after transfection. Results are the mean ± SEM of N = 3, n = 2. **H,I**. eATP (H) and LDH (I) from HEK-293T cells with a doxycycline–inducible GSDMD NT expression system treated with doxycycline (blue bars) or not (grey bars) for 3, 5 or 24 hours. Results are the mean ± SEM of N = 3, n = 2. **J.** Migration of THP-1 cells through a transwell towards supernatants from inducible HEK-293T cells treated as in H for 3 and 24 hours with doxycycline (dark blue) or not (light blue). In grey, the supernatants of doxycycline were treated with apyrase. Results are the mean + SEM for N = 3, n = 2. See also Figures S4, S5 and S6.

When we measured LDH levels in the same culture media, the resultant kinetics presented a gradual increase, reaching a maximum at 24 hours, the final time point measured (Fig. 3B). Notably, at 4 hours post-transfection, there was a significant release of ATP and less than 2% of LDH release. This early time point, where GSDMD-mediated cell permeabilization occurs before cell death, was selected to explore the possibility that different GSDMs isoforms could also facilitate ATP release.

Four hours after transfection, the expression of the different N-terminal fragments of GSDMA, B, C, and E resulted in significant ATP release compared to their full-length counterparts (Fig. 3C). At this time point, all GSDMs N-terminal fragments released less than 2% of LDH (Fig. 3D), showing that ATP can permeate through the various gasdermin pores before the onset of lytic cell death. As a control, we confirmed that after 24 hours of transfection, all GSDMs N-terminal fragments were able to induce lytic cell death, as evidenced by a significant increase of LDH release (Fig. S4).

Transfection of HEK-293T cells with a plasmid encoding NINJ1 led to an increase in both eATP concentration and LDH levels, which could be inhibited by glycine (Fig. 3E, F). This suggests that early cell lysis induced by NINJ1, even in the absence of GSDMs–membrane permeabilization, also triggers ATP release. In contrast, GSDMD N-terminal induced ATP release was independent on NINJ1, as glycine had no effect on this process (Fig. 3G). These findings collectively indicate that, although ATP is capable of being released through NINJ1 plasma membrane rupture as well as through GSDMs pores, the exit of ATP during NLRP3 induced pyroptosis is restricted to GSDMD pores and does not occur during the later phases of NINJ1–induced plasma membrane rupture.

### GSDMD–induced ATP released promotes monocyte migration

We finally investigated possible roles of GSDMD–dependent ATP release in immune cells using a recombinant doxycycline–induced system to control the formation of GSDMD pores. Using this system, we confirmed that ATP release increased after doxycycline addition, alongside a corresponding rise in LDH release (Fig. 3H, I). Similar to what we observed with transient transfections, the concentration of eATP in the supernatants decreased over time with doxycycline incubation, while LDH release progressively increased (Fig. 3H, I).

We then used these supernatants to assess their effect on THP-1 monocyte migration. Both, supernatants collected after 3 and 24 hours of doxycycline treatment stimulated THP-1 migration, with a significant greater number of monocytes migrating in response to the 3 hours supernatants (Fig. 3J, S5). Migration of THP-1 was inhibited when the supernatants were treated with apyrase (Fig. 3J), indicating that nucleotides released during GSDMD–induced cell permeabilization were responsible for stimulating THP-1 migration.

To test the ability of released ATP to reach neighboring cells, we added HEK-293T-GSDMD expressing full-length or N-terminal domain (2 hours after transient transfection) to a monolayer of HEK-293T cells stably expressing a chimeric plasma membrane luciferase (HEK-293T-pmeLUC) in the presence of luciferin. The time-course response detected by HEK-293T-pmeLUC cells showed that HEK-293T-GSDMD N-terminal domain released significantly more ATP than HEK-293T-GSDMD full-length, or mock transfected HEK-293T cells, and this ATP singling was prevented by treatment with apyrase (Fig. S6A). The HEK-293T-pmeLUC cells were calibrated with increasing concentrations of exogenous ATP (Fig. S6B) and the slope obtained (Fig. S6C) was used to calculate the concentration of ATP detected by HEK-293T-pmeLUC in each condition (Fig. S6A). The concentration of ATP released by HEK-293T-GSDMD N-terminal domain were in the mM range, showing that the paracrine extent of released ATP occurs at concentrations much higher than those detected in experiments in which ATP is measured in the entire supernatant.

Overall, our study highlights that ATP is released early during the initial permeabilization of the plasma membrane by GSDMD during NLRP3–induced pyroptosis, serving as an early and transient danger signal induces monocyte migration before the onset of lytic cell death.

## DISCUSSION

It is well established that eATP is a proinflammatory trigger that function as a second signal for the canonical NLRP3 inflammasome activation. In this report, we provide evidence that, in the early steps of inflammasome activation, ATP is released from macrophages in a GSDMD-dependent way. Our results suggest that upon NLRP3 inflammasome activation, ATP is transiently released and amplifies NLRP3 inflammasome effector responses by inducing monocyte migration.

Using 10^5^ BMDMs per 50 µl assay, the maximum observed eATP concentration was approximately 30 nM. Although this *in vitro* concentration seems relatively low for activating most P2 receptors, the high abundance of BMDMs in the bone marrow suggests that micromolar eATP concentrations could be expected, even when BMDMs displayed high eATP hydrolysis.

Moreover, low micromolar eATP is compatible with purinergic signaling, since eATP engage P2Y2, (EC_50_ ATP = 230 nM)^26^ to enhance IL-1β release following NLRP3 activation^27^. In recombinant systems expressing the GSDMD N-terminal fragment, eATP accumulates in micromolar levels. This supports the idea that GSDMD alone, without the upstream signaling that typically leads to its processing and membrane localization, can facilitate ATP release.

Moreover, it has been described that those cells can retain micromolar concentrations of eATP in their pericellular space, without significant nucleotide convection into the bulk milieu^28^. The delay in the YO-PRO-1 uptake and the reduced number of positive cells observed using P2X7-deficient macrophages also pointed in this direction. These experiments showed that the ATP released after NLRP3 activation could be enough to activate P2X7 receptor in the neighboring cells and also indicates that the YO-PRO-1 uptake could happen by different pathways at the same time: GSDMD and P2X7 receptor-associated mechanisms. In this context, the HEK-293T-pmeLUC system enabled the measurement of eATP in the immediate -nanometric-environment of the cell surface, where purinergic signaling occurs. Our experiments demonstrated that eATP released by HEK-293T-GSDMD NT was detected at high levels on the surface of HEK-293T-pmeLUC.

The relatively high eATP hydrolysis rate observed should also produce high concentrations of eADP, which in turn may activate P2Y1, P2Y12 and P2Y13 receptors^29^ and potentially trigger cellular responses with inflammatory relevancy. For example, the activation of P2Y12 receptor induces chemotaxis and chemokines release in tumor-associated macrophages^30^. In line, we found that extracellular nucleotides from cells expressing GSDMD pores were able to induce migration of THP-1, suggesting the attraction of cells towards pyroptotic cells. Although P2Y1 and P2Y13 ADP receptors were identified in macrophages^31^, their role in NLRP3 inflammasome or inflammation has not been characterized.

In addition to eADP accumulation caused by eATP hydrolysis, macrophages express ecto-nucleotidases NTPDase 1^32^ and 5’ NT^31^. The coupled action of both enzymes may produce adenosine from eATP, with consequent activation of P1 receptors. It is well known that the activation of P1 receptors by adenosine contributes to a broad range of anti-inflammatory responses, regulating inflammation^33^.

Therefore, the eATP kinetics after NLRP3 activation initiate an early pro-inflammatory phase characterized by the accumulation of eATP and its hydrolysis product, eADP. This is followed by a later phase where adenosine exerts anti-inflammatory effects, probably contributing to the fine-tuning of inflammation.

Our results using glycine, a cytoprotective amino acid that inhibits NINJ1 oligomerization and subsequent plasma membrane rupture, along with studies in *Ninj1*–deficient macrophages, showed that ATP is released via a non-lytic mechanism prior to pyroptotic cell death, as evidenced by ATP efflux occurring before LDH release. Notably, YO-PRO-1 uptake was not inhibited by glycine, indicating that this fluorescent molecule can pass through GSDMD pores before the final lytic phase of pyroptotic cell death. As already reported^16,22^, the sole ectopic expression of NINJ1 in cells induces lytic cell death independently of GSDM pores. Here, we further discovered that under these conditions NINJ1-oligomerization drives ATP release.

Our experiments showed that IL-1β and LDH release are distinct events. This aligns with previous studies showing that GSDMD pore formation triggers IL-1β (17 kDa) release independently of cell death^13,19,34^. In a subsequent phase of pyroptosis, GSDMD pores promotes NINJ1–mediated cell membrane rupture, leading to the release of LDH (a 140 kDa tetrameric complex), which is too large to pass through GSDMD pores^13,35^.

However, the mechanism by which GSDMD pores induce NINJ1 oligomerization and plasma membrane rupture remains unclear. Our findings, consistent with other studies, demonstrated that blocking NINJ1 oligomerization at the plasma membrane impaired LDH release, without affecting IL-1β release^,19,22,36^.

Noteworthy, eATP kinetics revealed a transient increase of eATP concentrations before the onset of lytic pyroptotic cell death, with no differences observed when NINJ1 was inhibited or genetically inactivated. This suggests that ATP release during pyroptosis is a controlled process. Moreover, we observed that intracellular concentrations of ATP dropped dramatically to less than 10% of their initial levels within 30 minutes of nigericin addition. LDH release begun around 25-30 minutes under our conditions, suggesting that cells are already in its path to silent death early after NLRP3 activation.

On the other hand, ATP and IL-1β release occur earlier than the lytic phase of pyroptotic cell death, suggesting that ATP release could be a general danger signal resulting from initial small GSDMD pores, which may be too small to allow for IL-1β release^37,38^. Our study further indicates that ATP release serves as a broad inflammatory danger signal, released through various GSDMs pores even in cells that do not express IL-1β, impacting both immune and non-immune cells. These findings indicate that ATP release through GSDMD pores is an early and transient non-lytic event occurring after the canonical NLRP3 inflammasome activation and before NINJ1– dependent cell lysis.

Limitations of the study: While our study provides significant insights into the early release of ATP mediated by GSDMD in BMDMs, the precise structural basis of ATP transport through GSDMD pores remains unresolved. While much progress has been made in understanding the molecular mechanisms regulating GSDMD pore formation—including activation by proteolytic cleavage, dynamics of pore assembly, post-translational modifications, and interplay with mitochondrial function—, how these regulatory mechanisms are coordinated in time and space, particularly in the context of ATP transmembrane permeability, remains to be further explored. Our analysis primarily focused on eATP as a danger signal, but other molecules (like e.g., other nucleotides) potentially released through GSDMD pores and their biological relevance were not investigated.

## Resource Availability

Data from this article is submitted as data source files accompanying the publication.

## Acknowledgments

We thank I. Couillin (CNRS Orleans, France) for *Gsdmd*^−/−^ mice, I. Hafner (National Institute of Chemistry, Ljubljana, Slovenia) for *Gsdmd*^−/−^ and *Ninj1*^−/−^ iBMDMs, F. Di Virgilio (University of Ferrara, Italy) for pmeLuc cells, E. Meunier (Institute of Pharmacology and Structural Biology, University of Toulouse, France) for inducible cells expressing GSDMD Nt, C. Martínez (IMIB, Murcia, Spain) for buffy coats, M.C. Baños and A.I. Gomez (IMIB, Murcia, Spain) for technical assistance with molecular and cellular biology and the members of the Pelegrin’s laboratory for comments and suggestions thought the development of this project.

## Author contributions

JS, DA-B, LH-N, AG, MP performed the experimental work and analyzed the data; JS, PJS analyzed data and performed regression analysis; MP, ABP-O, EM developed key stable cell lines; JS and PP, interpreted results, conceived the experiments, prepared the figures, paper writing and overall supervision of this study.

Conceptualization, Validation, Methodology, Data Curation and Writing – Original Draft: P.P. and J.S.; Formal Analysis: P.P., J.S., P.J.S., A.G., D.A.B., L.H.N., M.P.; Investigation: A.G., J.S., D.A.B., L.H.N.; Resources: M.P., A.B.P.O. and E.M.; Writing – Review and Editing: P.P., J.S. and P.J.S., Supervision and Project Administration: P.P.; Funding Acquisition: P.P. and J.S.

## Supplemental information

Document S1. Figures S1–S6

## Methods

### Animals

C57BL/6 (wild type, WT) mice were purchased from Harlan. P2X7 receptor-deficient mice (*P2rx7*^−/−^) were purchased from Jackson (Solle et al, 2001), and *Gsdmd*^−/−^, double caspase-1/11-deficient (*Casp1/11*^−/−^) (Heilig et al., 2018, Kuida et al., 1995) were all on C57BL/6 background. For all experiments, mice between 8–10 weeks of age bred under SPF conditions were used in accordance with the University Hospital Virgen Arrixaca animal experimentation guidelines, and the Spanish national (RD 53/2013 and Law 6/2013) and EU (86/609/EEC and 2010/63/ EU) legislation. According to legislation cited above, local ethics committee review approval is not needed, since mice were euthanized by CO_2_ inhalation and used to obtain bone marrow; no procedure was undertaken which compromised animal welfare (chapter 1, article 3 of RD 53/2013).

### Cells and treatments

Bone marrow derived-macrophages (BMDMs) were obtained from wild type or knockout mice as described (Barbera-Cremades et al., 2012) and maintained in Dulbecco’s modified Eagle’s medium (DMEM-F12) supplemented with 10 % FBS and 2 mM GlutaMax and 1% of penicilin-streptomycin.

Blood samples from healthy humans volunteers included in this study, who gave written informed consent, were collected, and processed following standard operating procedures with appropriate approval of the Ethical Committee of the Clinical University Hospital Virgen de la Arrixaca (Murcia, Spain) with reference #2021-7-9-HCUVA. Human peripheral blood mononuclear cells (PBMCs) were isolated from whole blood samples using Ficoll-based gradient separation method. The blood samples were added on the top chamber of a SepMate Falcon tube, containing 15 mL of Ficoll with 1.077 g/mL density. The tubes were centrifugated at 1.200 x g during 10 min and the PBMCs fraction was removed. The PMBCs were washed with PBS, suspended on OptiMEM, counted and 2 x 10^8^ cell were added to 10 mL of Percoll. The tubes were centrifugated at 580 x g and the monocytes obtained were suspended in RPMI 1640 medium supplemented with 10 % of human serum and seeded on 24-well plates at a density of 5×10^5^ cells by well. After 3 days, the cells were washed and the differentiated macrophages were maintained 4 days until use.

Immortalized bone marrow-derived macrophages (iBMDMs) WT, *Gsdmd*^−/−^ and *Ninj1*^−/−^ were a gift from Dr. I. Hafner (National Institute of Chemistry, Ljubljana, Slovenia). The iBMDMs were grown in DMEM-F12 supplemented with 10% FBS.

THP-1 cells were maintained in RPMI 1640 media supplemented with 10% FBS and to measure ATP release, THP-1 were differentiated to macrophages with 0.5 µM PMA for 30 min before NLRP3 activation. For cell migration, THP-1 were not treated with PMA.

All types of macrophages were primed with LPS (1 µg/mL, 4h) and subsequent canonical NLRP3 inflammasome activation was achieved with nigericin at the indicated concentrations.

HEK-293T cells were maintained in DMEM-F12 supplemented with 10 % FBS. For transfection with GSDMA, GSDMB, GSDMC, GSDMD, GSDME and NINJ1, cells were incubated at least 16 to allow adhesion to the plate before cationic lipid-based transfection using Lipofectamine 2000 (Invitrogen) according to the manufacturer’s instruction. Briefly, two solutions of 50 µl of Opti-MEM containing a mix of 1-2 µg of total plasmid DNA (tube A) and 3 µl of Lipofectamine 2000 (tube B) were prepared and incubated 5 min at room temperature. After this period of time, the volume of DNA tube (A) was added into Lipofectamine 2000 tube (B), mixed gently and incubated 20 min at RT to allow lipid/DNA complexes formation and then added drop by drop to the cells. The plate was swirled gently and incubated at 37 °C until the times indicated to extracellular ATP or LDH determinations.

HEK-293T stably transfected with plasma membrane luciferase (HEK-293T-pmeLUC)^39^ were a gift of Prof. F. Di Virgilio (University of Ferrara, Italy). Cells were maintained in DMEM-F12 supplemented with 10 % FBS and 1% of penicillin-streptomycin.

### Generation of HEK-293T with an inducible GSDMD NT system

HEK293FT TREx-GSDMD-NT inducible system by doxycycline were generated by T-REx™ Complete Kit, with pcDNA™4/TO Vector (K102001, ThermoFisher). The cells were maintained in DMEM-F12 supplemented with 20 % FBS, 1% L-Glutamine, 1% of penicillin-streptomycin (Corning) and with 100µg/ml zeocin (Gibco). Induction of GSDMD-NT was achieved by doxycycline treatment (10ng/ml) for 3, 5 or 24 hours.

### Extracellular ATP measurements

Extracellular ATP concentration in the supernatants were measured with the luciferase-luciferin assay (Strehler, 2006). BMDMs were plated in a 96-well plate and the culture media was replaced with basal salt solution containing 130 mM NaCl, 5 mM KCl, 1.5 mM CaCl_2_, 1 mM MgCl_2_, 25 mM Na-HEPES (adjusted to pH 7.5 at room temperature), 5 mM glucose, 0.1 % BSA, 47.8 µM D-Luciferin, 0,5 µM luciferase and 0.1 mg/mL of CoA. For extracellular ATP calculations, a standard curve (from 10 to 200 nM ATP) was used. Bioluminescence measured were performed in a BioTek Synergy Neo2 Hybrid Multimode Reader at room temperature, because the luciferase activity at 37°C is only 10% of that observed at 20°C (Gorman et al., 2003).

### YO-PRO-1 uptake assay

YO-PRO-1 uptake was measured by fluorescence microscopy. BMDMs were seeded on coverslips and before the experiment were washed two times with basal salt solution. After that, 2 µM YO-PRO-1 was added to the assay media (saline solution) and the macrophages were treated with nigericin 25 µM. Images were acquired every 30 secs at room temperature with a Nikon Eclipse Ti microscope using a 20x S Plan Fluor objective (numerical aperture 0.45), a digital Sight DS-QiMc camera (Nikon), 482nm/536nm filter set (Semrock) and NIS Elements software (Nikon). The fluorescence intensity related to basal were analyzed using Image J software (US National Institute of Health). Alternatively, YO-PRO-1 uptake was measured at room temperature using a Synergy Neo2 Hybrid Multimode plate reader (BioTek).

### ELISA

IL-1β release was measured by ELISA for mouse IL-1β (R&D Systems) following the manufacturer’s instructions and read in a Synergy Mx plate reader (BioTek). The concentration of IL-1β was estimated using a standard with known concentrations of recombinant IL-1β.

### Lactate dehydrogenase (LDH) assay

LDH release was measured using the Cytotoxicity Detection kit (Roche) following the manufacturer’s instructions, and expressed as percentage of total cell LDH content.

### Monocyte migration assay

HEK-293T cells with an inducible system for GSDMD NT were seeded on 24-well plates at a density of 3×10^5^ cells by well. The next day the cells were incubated with 10 ng/mL doxycycline for 3 and 24 hours in OptiMEM media. After this time, the supernatants were collected, centrifugated at 16,060 x*g* for 30 sec and 500 µL of the free-cells supernatants were loaded into the lower chamber of a transwell insert (Sarstedt 83.3932.500_PET 5µm) placed on 24-well plate. At this point, 40 U/ml apyrase (Sigma-Aldrich) was added to the media of the well. In addition, the supernatant of HEK-293T cells with an inducible system for GSDMD NT not treated with doxycycline was supplemented with 10 ng/ml doxycycline to ensure that THP-1 chemotaxis is not due to the presence of doxycycline. After 30 min at 37°C, 3×10^5^ THP-1 cells were loaded into the upper chamber of the transwell inserts, and THP-1 cells were supplemented with 10 ng/ml doxycycline again to corroborate that the presence of doxycycline do not induce migration. After 3 or 24 hours of incubation at 37°C, the inserts were removed and 16,000 units of flow cytometry counting beads (BD Trucount Tubes) were added per well. The migrated THP-1 with the counting beads were transferred to flow cytometry tubes. The samples were analyzed by flow cytometry (LSR Fortessa flow citometer, BD) recording events by gating on beads and collecting a fixed number of bead events per sample (730) and analyzed in FCS Express 5 Flow cytometry software (De novo Software). The percent of migration were calculated compared with total cells.

### Quantification and statistical analysis

#### Statistical methods

The data are presented as the mean ± SEM. Data were analyzed using Prism Version 9.5.1 (GraphPad) software the non-parametric Mann-Whitney test (two tails, 95% confidence level), *p<0.05; ns, not significant (p>0.05) difference.

## SUPPLEMENTARY FIGURES

**Figure S1.**
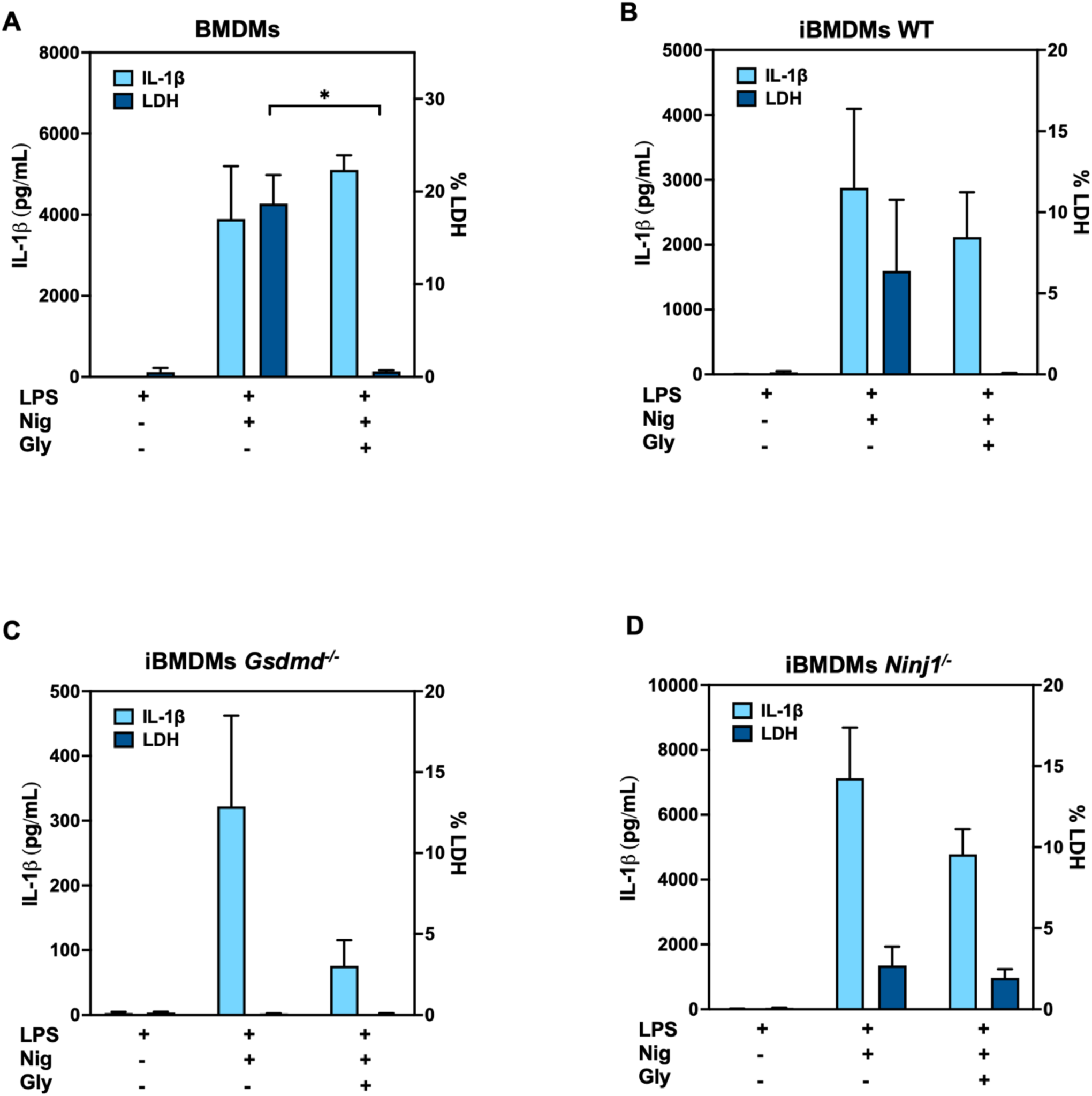
Glycine impairs LDH but not IL-1β release after inflammasome activation. **A-D.** IL-1β (light blue bars) and LDH (dark blue bars) release from LPS-primed macrophages, treated with 20 µM nigericin (Nig) with or without 5 mM glycine (Gly) for 1 hour. The supernatants from bone marrow derived macrophages (BMDMs) is shown in A, from immortalized BMDMs (iBMDMs) WT in B, iBMDMs *Gsdmd*^−/−^ in C, and iBMDMs *Ninj1*^−/−^ in D. Data are means + SEM of independent experiments (N) performed in duplicate (n), N= 4, n = 2. Related to Figure 1.

**Figure S2.**
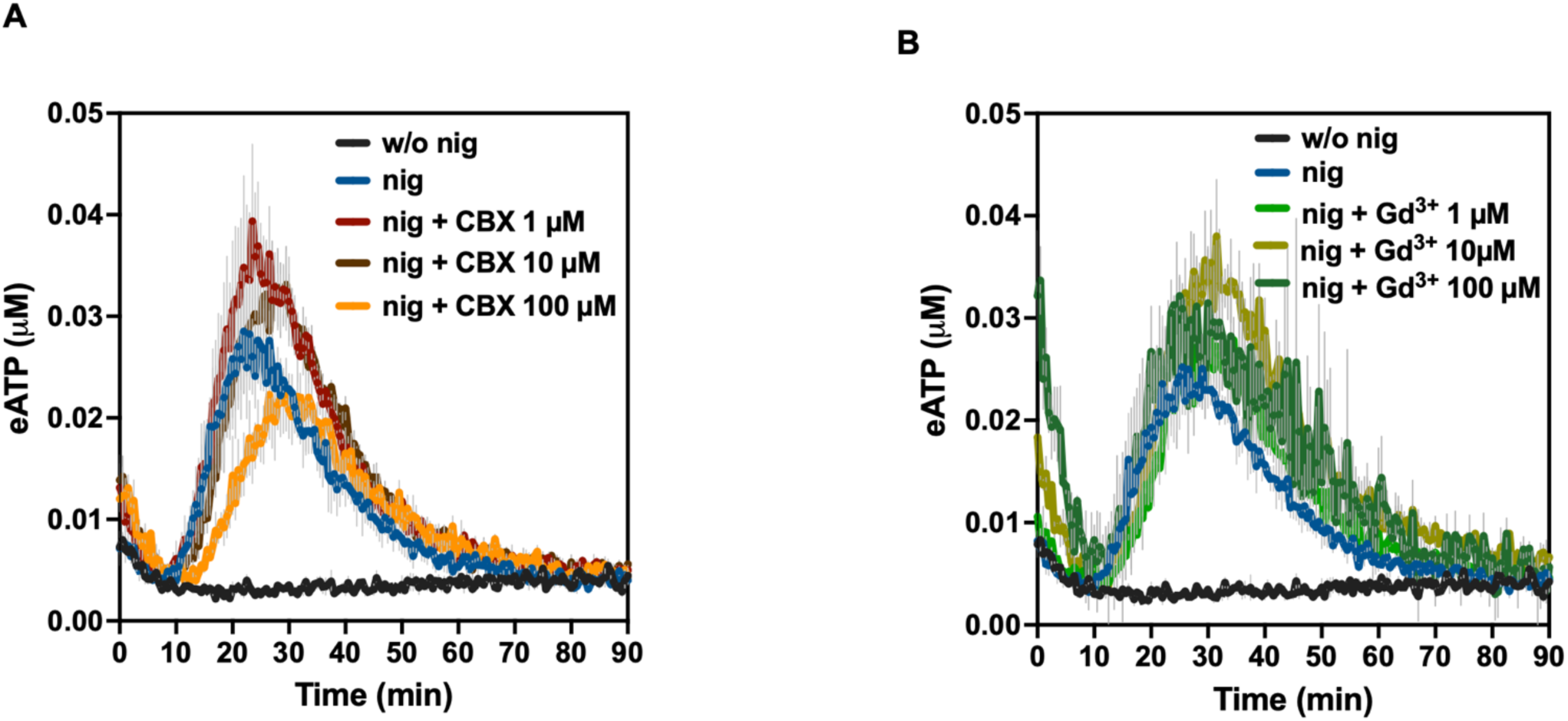
ATP release after NLRP3 inflammasome activation is not dependent on pannexin-1 or connexins. **A,B**. LPS-primed bone marrow derived macrophages (BMDMs) treated with saline solution (black lines), nigericin (nig) 25 µM (blue lines) in the presence of 1, 10 and 100 µM carbenoxolone (CBX) (A), or 1, 10 and 100 µM gadolinium (Gd^3+^) (B). Data are means ± SEM of independent experiments (N) performed in duplicate (n), N= 4, n = 2. Related to Figure 1.

**Figure S3.**
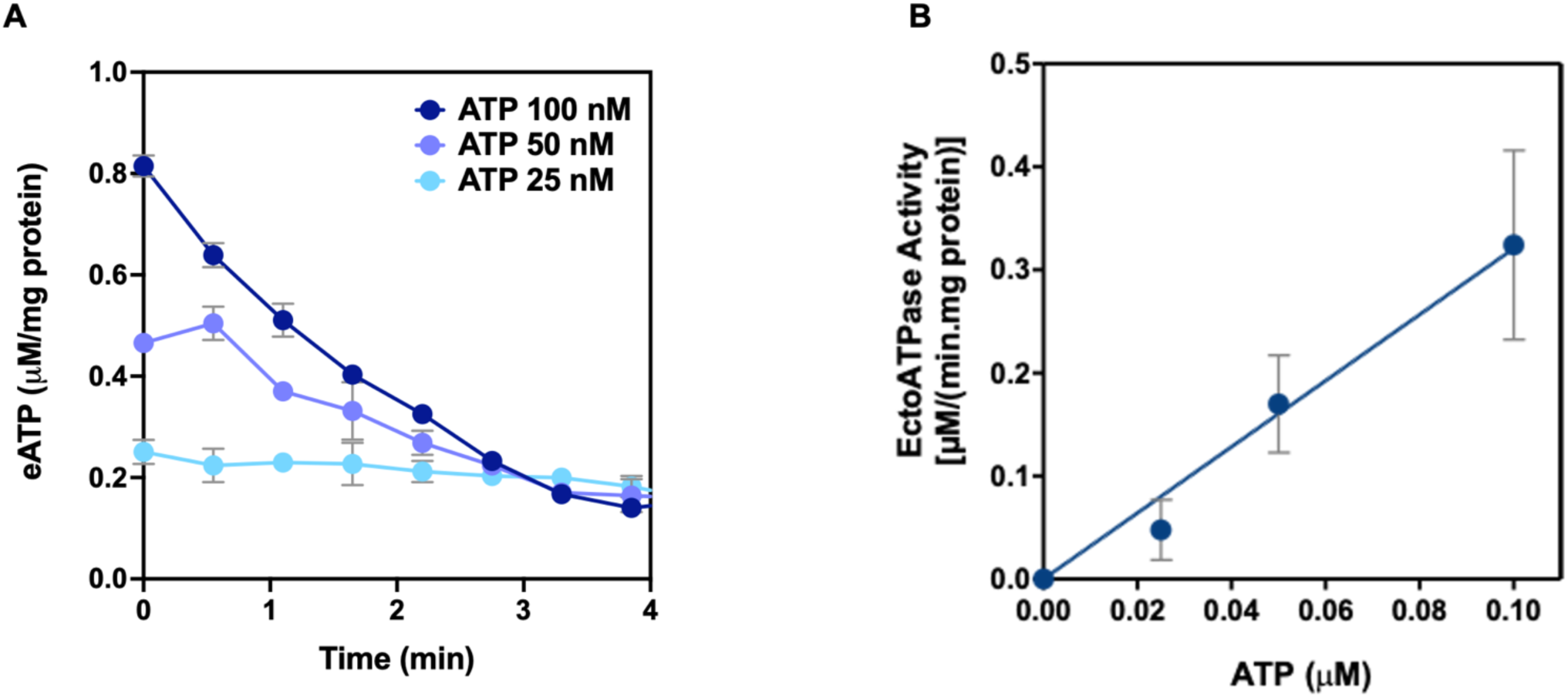
Degradation of eATP on the surface of macrophages. **A.** Kinetic of exogenous ATP decay added to LPS-primed bone marrow derived macrophages (BMDMs) activated with 25 μM nigericin and incubated with ATP (25 nM, light blue line; 50 nM, blue line, and 100 nM, dark blue line) for 4 minutes. **B.** Substrate curve of eATP hydrolysis built with the data obtained from A. Results are means ± SEM, and are expressed as µM ATP/(min.mg protein). Data are means ± SEM of independent experiments (N) performed in duplicate (n), N = 3, n = 2. The continuous line represents fitting of a linear function to the experimental data with a slope = K_ATP_. Related to Figure 1.

**Figure S4.**
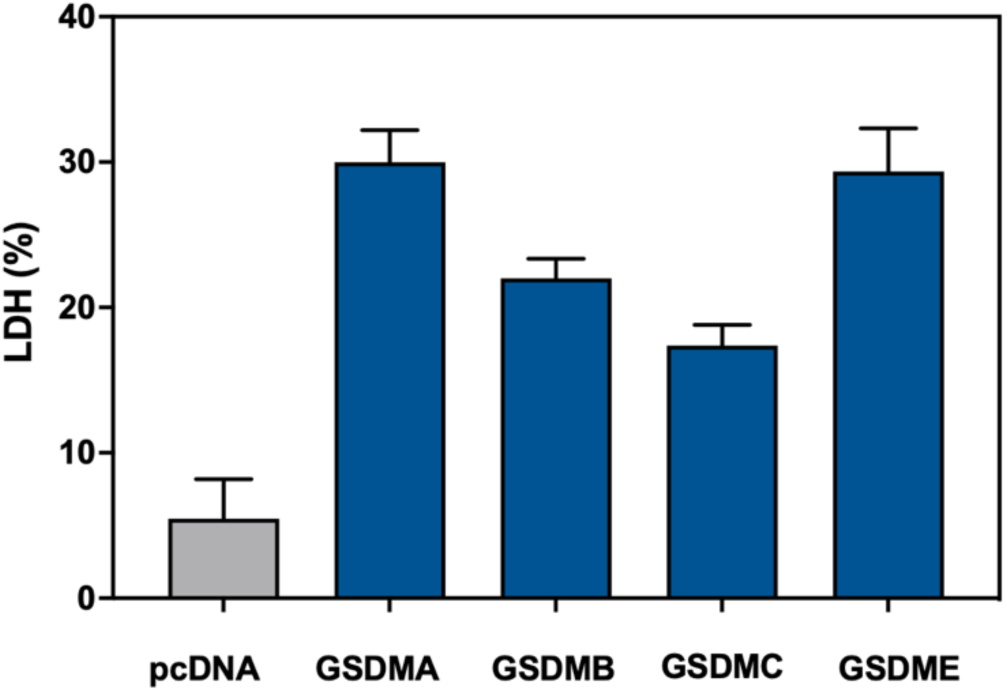
LDH release after expression of GSDMs in HEK-293T. Extracellular LDH (percentage to total intracellular LDH) from HEK-293T cells transfected with pcDNA3.1 (grey bar), or the N-terminal fragments of GSDMA, GSDMB, GSDMC and GSDME (blue bars) for 24 hours. Data are means ± SEM of independent experiments (N) performed in duplicate (n), N = 3, n = 2. Related to Figure 3.

**Figure S5.**
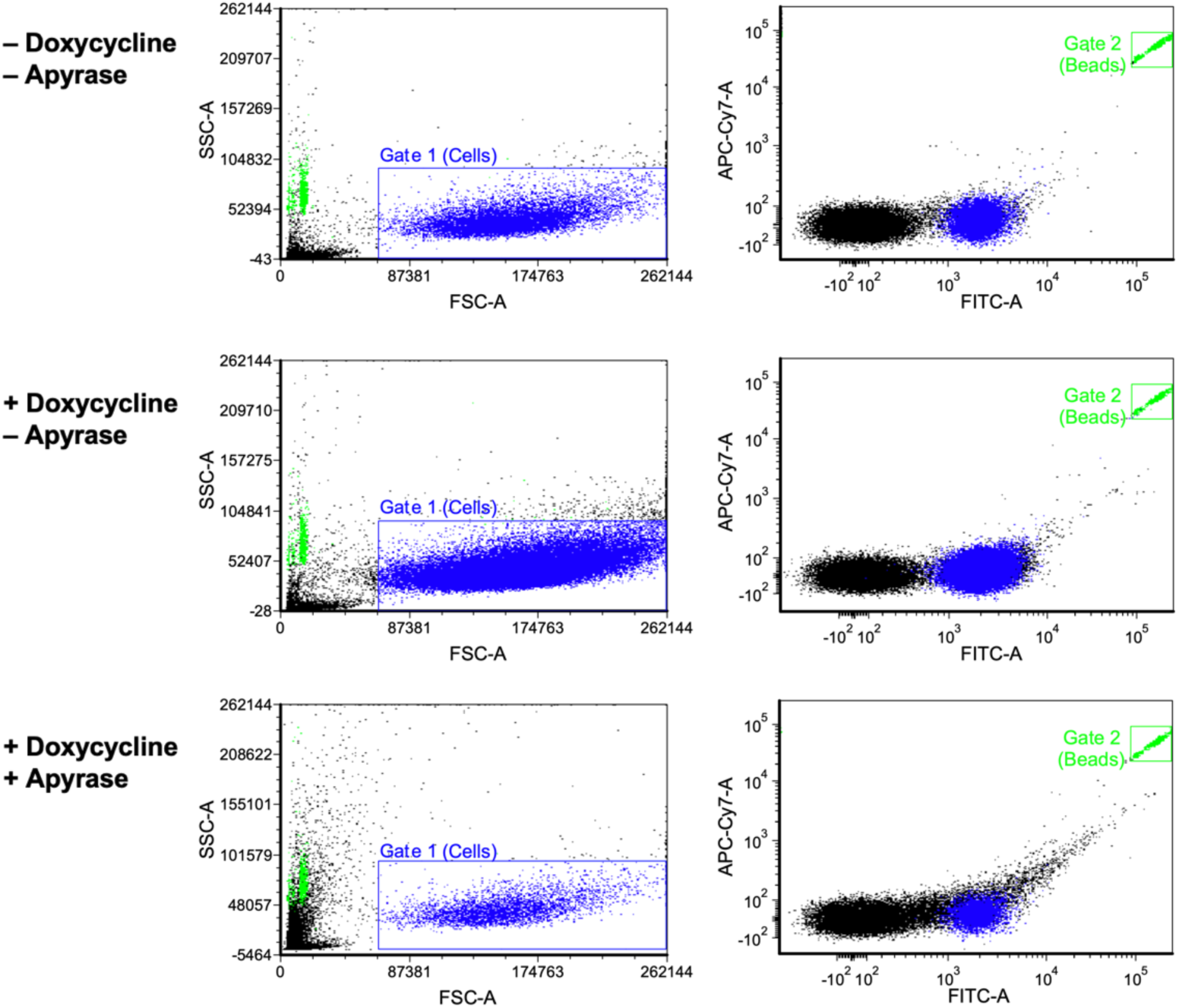
Representative gating strategy for THP-1 migration. Gating strategy followed to detect migration of THP-1 monocytes (Gate 1, cells in blue color) through a transwell to supernatants from HEK-293T-GSDMD NT treated or not with doxycycline for 3 hours. Apyrase was added to a supernatant of doxycycline treated cells. Flow cytometry calibration beads are shown in Gate 2 (green color). One representative experiment out of N=3 is shown. Related to Figure 3.

**Figure S6.**
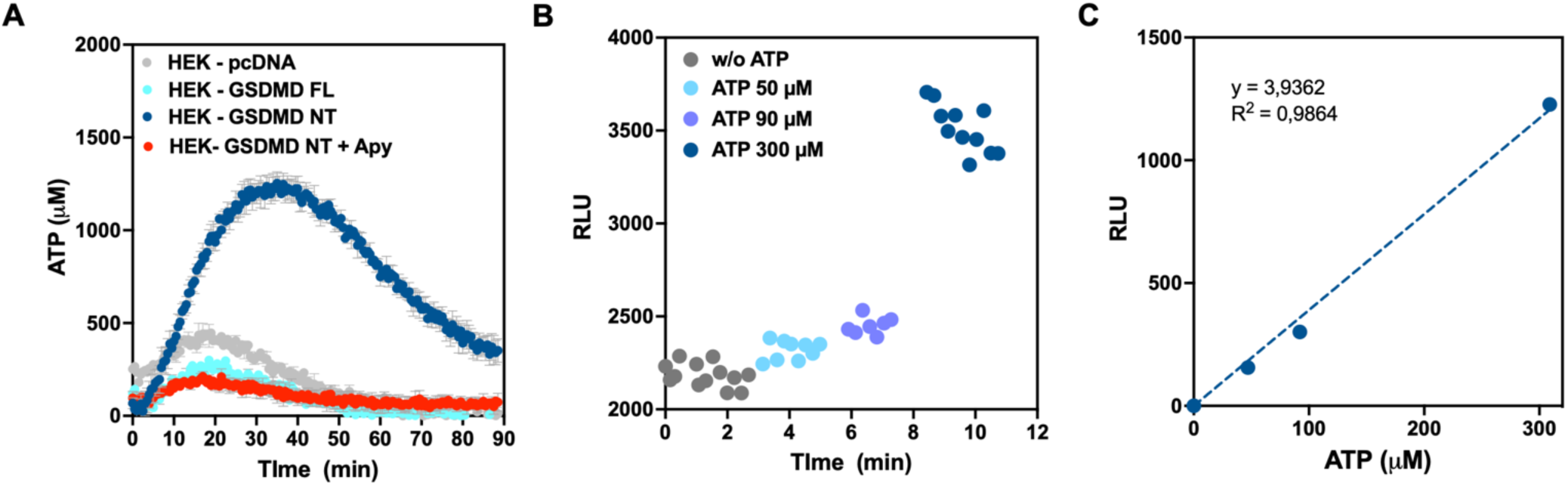
HEK-293T-pmeLUC cells detect ATP release from HEK-293T cells expressing GSDMD NT. Relative units of luminescence (RLUs) emitted by HEK-293T cells stably expressing pmeLUC construction after addition of HEK-293T cells transiently transfected for 2 hours with pcDNA3.1 (grey), GSDMD FL (light blue), or GSDMD NT (dark blue) in the presence of 200 µM luciferin and with or without 20 U/mL apyrase (red). Data are means ± SEM of one representative experiment performed in triplicate. N = 2. **A.** Relative units of luminescence (RLUs) emitted by HEK-293T cells stably expressing pmeLUC construction after addition of 50, 90 and 300 µM ATP in the presence of 200 µM luciferin. **B.** Dose-response fitting from data obtained in A. The slope of the linear regression was used to transform the RLU into ATP concentrations in A. Related to Figure 3.

## REFERENCES

1. Giuliani, A.L., Sarti, A.C., and Di Virgilio, F. (2019). Extracellular nucleotides and nucleosides as signalling molecules. Immunol. Lett. 205, 16–24. 10.1016/J.IMLET.2018.11.006.

2. Cekic, C., and Linden, J. (2016). Purinergic regulation of the immune system. Nat. Rev. Immunol. 16, 177–192. 10.1038/NRI.2016.4.

3. Kepp, O., Loos, F., Liu, P., and Kroemer, G. (2017). Extracellular nucleosides and nucleotides as immunomodulators. Immunol. Rev. 280, 83–92. 10.1111/IMR.12571.

4. Kobayashi, D., Umemoto, E., and Miyasaka, M. (2023). The role of extracellular ATP in homeostatic immune cell migration. Curr. Opin. Pharmacol. 68, 102331. 10.1016/J.COPH.2022.102331.

5. Fitz, J.G. (2007). Regulation of cellular ATP release. Trans. Am. Clin. Climatol. Assoc. 118, 199–208.

6. Dosch, M., Gerber, J., Jebbawi, F., and Beldi, G. (2018). Mechanisms of ATP Release by Inflammatory Cells. Int. J. Mol. Sci. 19. 10.3390/IJMS19041222.

7. Taruno, A. (2018). ATP Release Channels. Int. J. Mol. Sci. 19. 10.3390/IJMS19030808.

8. Zimmermann, H. (2021). Ectonucleoside triphosphate diphosphohydrolases and ecto-5’-nucleotidase in purinergic signaling: how the field developed and where we are now. Purinergic Signal. 17, 117–125. 10.1007/S11302-020-09755-6.

9. Zimmermann, H. (2021). History of ectonucleotidases and their role in purinergic signaling. Biochem. Pharmacol. 187, 114322. 10.1016/j.bcp.2020.114322.

10. Pelegrin, P. (2021). P2X7 receptor and the NLRP3 inflammasome: Partners in crime. Biochem. Pharmacol. 187. 10.1016/J.BCP.2020.114385.

11. Tapia-Abellán, A., Angosto-Bazarra, D., Alarcón-Vila, C., Baños, M.C., Hafner-Bratkovič, I., Oliva, B., and Pelegrín, P. (2021). Sensing low intracellular potassium by NLRP3 results in a stable open structure that promotes inflammasome activation. Sci. Adv. 7. 10.1126/SCIADV.ABF4468.

12. Liu, X., Zhang, Z., Ruan, J., Pan, Y., Magupalli, V.G., Wu, H., and Lieberman, J. (2016). Inflammasome-activated gasdermin D causes pyroptosis by forming membrane pores. Nature 535, 153–158. 10.1038/NATURE18629.

13. Evavold, C.L., Ruan, J., Tan, Y., Xia, S., Wu, H., and Kagan, J.C. (2018). The Pore-Forming Protein Gasdermin D Regulates Interleukin-1 Secretion from Living Macrophages. Immunity 48, 35–44.e6. 10.1016/J.IMMUNI.2017.11.013.

14. Heilig, R., Dick, M.S., Sborgi, L., Meunier, E., Hiller, S., and Broz, P. (2018). The Gasdermin-D pore acts as a conduit for IL-1β secretion in mice. Eur. J. Immunol. 48, 584–592. 10.1002/EJI.201747404.

15. Kelley, N., Jeltema, D., Duan, Y., and He, Y. (2019). The NLRP3 Inflammasome: An Overview of Mechanisms of Activation and Regulation. Int. J. Mol. Sci. 20. 10.3390/IJMS20133328.

16. Kayagaki, N., Kornfeld, O.S., Lee, B.L., Stowe, I.B., O’Rourke, K., Li, Q., Sandoval, W., Yan, D., Kang, J., Xu, M., et al. (2021). NINJ1 mediates plasma membrane rupture during lytic cell death. Nature 591, 131–136. 10.1038/S41586-021-03218-7.

17. Degen, M., Santos, J.C., Pluhackova, K., Cebrero, G., Ramos, S., Jankevicius, G., Hartenian, E., Guillerm, U., Mari, S.A., Kohl, B., et al. (2023). Structural basis of NINJ1-mediated plasma membrane rupture in cell death. Nature 618, 1065–1071. 10.1038/S41586-023-05991-Z.

18. Baroja-Mazo, A., Martín-Sánchez, F., Gomez, A.I., Martínez, C.M., Amores-Iniesta, J., Compan, V., Barberà-Cremades, M., Yagüe, J., Ruiz-Ortiz, E., Antón, J., et al. (2014). The NLRP3 inflammasome is released as a particulate danger signal that amplifies the inflammatory response. Nat. Immunol. 15, 738–748. 10.1038/NI.2919.

19. de Torre-Minguela, C., Gómez, A.I., Couillin, I., and Pelegrín, P. (2021). Gasdermins mediate cellular release of mitochondrial DNA during pyroptosis and apoptosis. FASEB J. 35. 10.1096/FJ.202100085R.

20. Phulphagar, K., Kühn, L.I., Ebner, S., Frauenstein, A., Swietlik, J.J., Rieckmann, J., and Meissner, F. (2021). Proteomics reveals distinct mechanisms regulating the release of cytokines and alarmins during pyroptosis. Cell Rep. 34. 10.1016/J.CELREP.2021.108826.

21. Fink, S.L., and Cookson, B.T. (2006). Caspase-1-dependent pore formation during pyroptosis leads to osmotic lysis of infected host macrophages. J. Immunol. 202, 1913– 1926. 10.1111/J.1462-5822.2006.00751.X.

22. Borges, J.P., Sætra, R.S.R., Volchuk, A., Bugge, M., Devant, P., Sporsheim, B., Kilburn, B.R., Evavold, C.L., Kagan, J.C., Goldenberg, N.M., et al. (2022). Glycine inhibits NINJ1 membrane clustering to suppress plasma membrane rupture in cell death. Elife 11. 10.7554/ELIFE.78609.

23. Muñoz, M.F., Griffith, T.N., and Contreras, J.E. (2021). Mechanisms of ATP release in pain: role of pannexin and connexin channels. Purinergic Signal. 17, 549–561. 10.1007/S11302-021-09822-6.

24. Schachter, J., Alvarez, C.L., Bazzi, Z., Faillace, M.P., Corradi, G., Hattab, C., Rinaldi, D.E., Gonzalez-Lebrero, R., Molineris, M.P., Sévigny, J., et al. (2021). Extracellular ATP hydrolysis in Caco-2 human intestinal cell line. Biochim. Biophys. acta. Biomembr. 1863. 10.1016/J.BBAMEM.2021.183679.

25. Rühl, S., Shkarina, K., Demarco, B., Heilig, R., Santos, J.C., and Broz, P. (2018). ESCRT-dependent membrane repair negatively regulates pyroptosis downstream of GSDMD activation. Science 362, 956–960. 10.1126/SCIENCE.AAR7607.

26. Jacobson, K.A., Delicado, E.G., Gachet, C., Kennedy, C., von Kügelgen, I., Li, B., Miras-Portugal, M.T., Novak, I., Schöneberg, T., Perez-Sen, R., et al. (2020). Update of P2Y receptor pharmacology: IUPHAR Review 27. Br. J. Pharmacol. 177, 2413–2433. 10.1111/BPH.15005.

27. de la Rosa, G., Gómez, A.I., Baños, M.C., and Pelegrín, P. (2020). Signaling Through Purinergic Receptor P2Y2 Enhances Macrophage IL-1β Production. Int. J. Mol. Sci. 2020, Vol. 21, Page 4686 21, 4686. 10.3390/IJMS21134686.

28. Yegutkin, G.G., Mikhailov, A., Samburski, S.S., and Jalkanen, S. (2006). The Detection of Micromolar Pericellular ATP Pool on Lymphocyte Surface by Using Lymphoid Ecto-Adenylate Kinase as Intrinsic ATP Sensor. Mol. Biol. Cell 17, 3378. 10.1091/MBC.E05-10-0993.

29. von Kügelgen, I. (2021). Molecular pharmacology of P2Y receptor subtypes. Biochem. Pharmacol. 187. 10.1016/J.BCP.2020.114361.

30. Kloss, L., Dollt, C., Schledzewski, K., Krewer, A., Melchers, S., Manta, C., Sticht, C., Torre, C. de la, Utikal, J., Umansky, V., et al. (2019). ADP secreted by dying melanoma cells mediates chemotaxis and chemokine secretion of macrophages via the purinergic receptor P2Y12. Cell Death Dis. 10. 10.1038/S41419-019-2010-6.

31. Merz, J., Nettesheim, A., von Garlen, S., Albrecht, P., Saller, B.S., Engelmann, J., Hertle, L., Schäfer, I., Dimanski, D., König, S., et al. (2021). Pro- and anti-inflammatory macrophages express a sub-type specific purinergic receptor profile. Purinergic Signal. 17, 481–492. 10.1007/S11302-021-09798-3.

32. Lévesque, S.A., Kukulski, F., Enjyoji, K., Robson, S.C., and Sévigny, J. (2010). NTPDase1 governs P2X7-dependent functions in murine macrophages. Eur. J. Immunol. 40, 1473–1485. 10.1002/EJI.200939741.

33. Antonioli, L., Fornai, M., Blandizzi, C., Pacher, P., and Haskó, G. (2019). Adenosine signaling and the immune system: When a lot could be too much. Immunol. Lett. 205, 9–15. 10.1016/J.IMLET.2018.04.006.

34. Xia, S., Zhang, Z., Magupalli, V.G., Pablo, J.L., Dong, Y., Vora, S.M., Wang, L., Fu, T.M., Jacobson, M.P., Greka, A., et al. (2021). Gasdermin D pore structure reveals preferential release of mature interleukin-1. Nature 593, 607–611. 10.1038/S41586-021-03478-3.

35. Russo, H.M., Rathkey, J., Boyd-Tressler, A., Katsnelson, M.A., Abbott, D.W., and Dubyak, G.R. (2016). Active Caspase-1 Induces Plasma Membrane Pores That Precede Pyroptotic Lysis and Are Blocked by Lanthanides. J. Immunol. 197, 1353–1367. 10.4049/JIMMUNOL.1600699.

36. Verhoef, P.A., Kertesy, S.B., Lundberg, K., Kahlenberg, J.M., and Dubyak, G.R. (2005). Inhibitory effects of chloride on the activation of caspase-1, IL-1beta secretion, and cytolysis by the P2X7 receptor. J. Immunol. 175, 7623–7634. 10.4049/JIMMUNOL.175.11.7623.

37. Santa Cruz Garcia, A.B., Schnur, K.P., Malik, A.B., and Mo, G.C.H. (2022). Gasdermin D pores are dynamically regulated by local phosphoinositide circuitry. Nat. Commun. 13. 10.1038/S41467-021-27692-9.

38. Tsuchiya, K., Nakajima, S., Hosojima, S., Thi Nguyen, D., Hattori, T., Manh Le, T., Hori, O., Mahib, M.R., Yamaguchi, Y., Miura, M., et al. (2019). Caspase-1 initiates apoptosis in the absence of gasdermin D. Nat. Commun. 10. 10.1038/S41467-019-09753-2.

39. Pellegatti, P., Falzoni, S., Pinton, P., Rizzuto, R., and Di Virgilio, F. (2005). A novel recombinant plasma membrane-targeted luciferase reveals a new pathway for ATP secretion. Mol. Biol. Cell 16, 3659–3665. 10.1091/MBC.E05-03-0222.

